# Microsaccades transiently lateralise EEG alpha activity

**DOI:** 10.1101/2022.09.02.506318

**Authors:** Baiwei Liu, Anna C. Nobre, Freek van Ede

## Abstract

The lateralisation of 8-12 Hz alpha activity is a canonical signature of human spatial cognition that is typically studied under strict fixation requirements. Yet, even during attempted fixation, the brain produces small involuntary eye movements known as microsaccades. Here we report how spontaneous microsaccades – made in the absence of incentives to look elsewhere – can themselves drive transient lateralisation of EEG alpha power according to microsaccade direction. This transient lateralisation of posterior alpha power occurs similarly following start and return microsaccades and is driven by increased alpha power ipsilateral to microsaccade direction. This reveals new links between spontaneous microsaccades and human electrophysiological brain activity, and highlights how microsaccades are an important factor to consider in studies relating alpha activity to spatial cognition.

## INTRODUCTION

In laboratory investigations into the neural basis of vision and cognition, it is common to instruct participants to maintain fixation. Ensuring control over where visual inputs land on the retina helps situate inputs relative to particular brain areas or receptive fields. Yet, we know from ample research that true fixation is an illusion ^1^. Even during attempted fixation, the brain generates small involuntary eye movements, commonly referred to as microsaccades ^1–3^. Microsaccades, and their retinal consequences, are themselves associated with specific neural modulations. However, to date, systematic investigations into the neural consequences of microsaccades have taken place predominantly at the level of single-cell recording in non-human primates, as reviewed in ^4,5^.

It is paramount to characterise the neural consequences of microsaccades also in humans. Not only may these consequences be of interest in and of themselves, but such consequences may also account for observed neural modulations by specific cognitive task variables when these variables also modulate microsaccades. This may not be an uncommon scenario given that microsaccade direction and rate can be modulated by spatial ^6–10^ and temporal ^11–13^ cognitive variables. One particularly well-established example is the directional biasing of microsaccades by the ‘covert’ allocation of spatial attention to peripheral locations ^6–10^.

While several prior studies have investigated how microsaccades modulate neural activity in humans^10,14–18^, key questions remain unexplored. In particular, it has remained unaddressed whether the direction of microsaccades is associated with specific spatial modulations of EEG activity in the classic 8-12 Hz alpha band. The underexplored nature of the link between microsaccade direction and spatial modulation of EEG-alpha activity is surprising provided that alpha lateralisation is a canonical signature of spatial cognition in human cognitive neuroscience ^19–24^. Here we fill this gap and report how spontaneous microsaccades – made in the absence of incentives to look elsewhere – are associated with transient lateralisation of EEG activity in the alpha frequency band according to microsaccade direction.

## RESULTS

We investigated microsaccades during the delay period of a working-memory task when only a fixation marker remained on the screen, which participants were instructed to look at. Despite this instruction, we identified numerous microsaccades during the delay **(Supplementary Fig. 1)**. Because there was no incentive to look at places other than fixation during the delay, we will refer to these microsaccades as “spontaneous”. Because we observed a similar proportion of microsaccades to the left and right **(Supplementary Fig. 1)**, we could investigate the lateralisation of EEG-alpha activity according to microsaccade direction. To do so, we aligned our EEG data to microsaccade onset (identified using gaze velocity; **Fig. 1a**) and compared activity in posterior EEG electrodes contralateral vs. ipsilateral to the direction of the identified spontaneous microsaccades.

**Figure 1.**
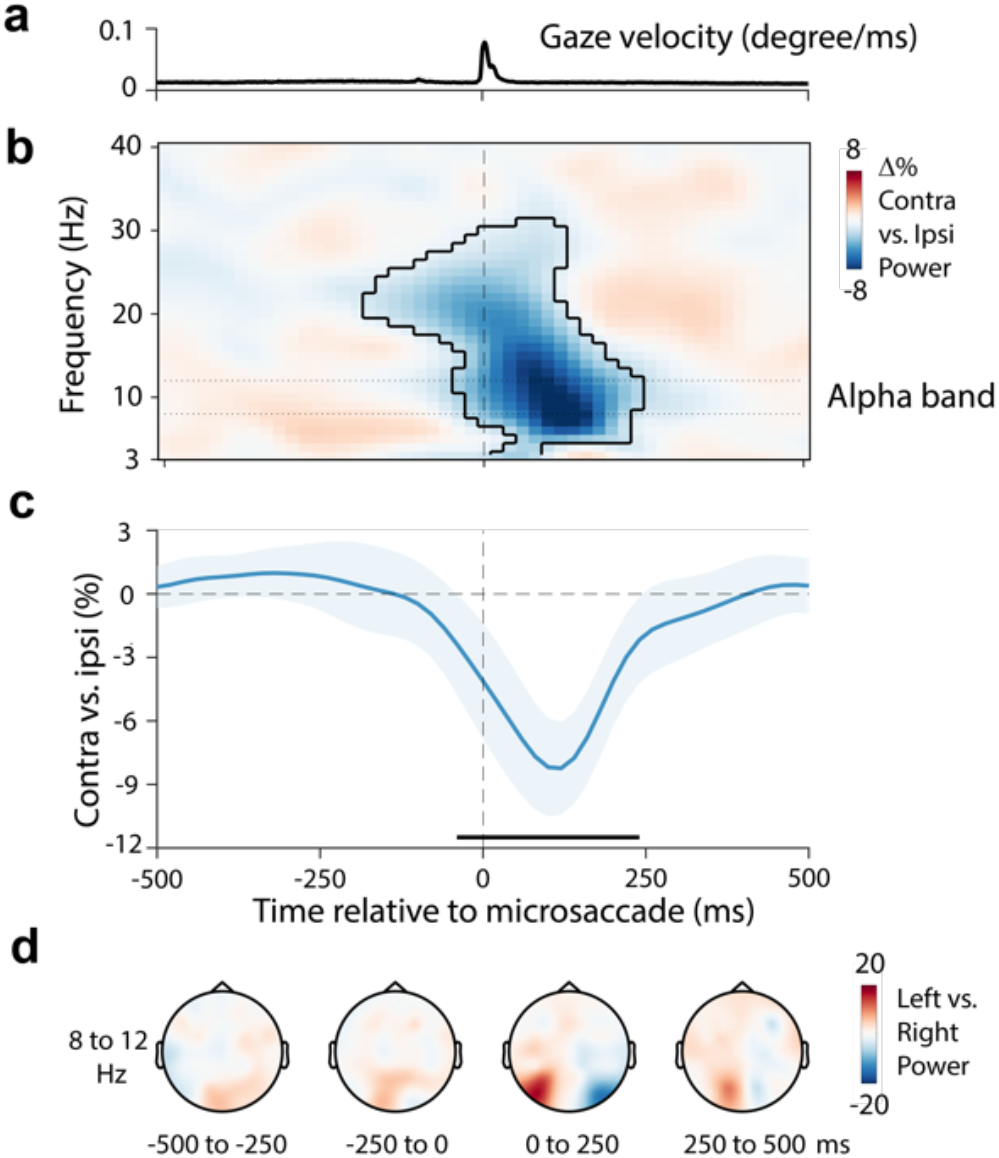
Microsaccades transiently lateralise EEG alpha power. **a)** Time course of gaze velocity. **b)** Spectral lateralisation (contralateral minus ipsilateral) in selected posterior electrodes (PO7, PO8) relative to the left/right microsaccade direction. Outline indicates the significant time-frequency cluster (two-sided cluster-based permutation test). **c)** Time course of 8-12 Hz lateralisation. The black horizontal line indicates the significant temporal cluster (two-sided cluster-based permutation test). **d)** Topographical maps of the difference between 8-12 Hz power for left vs. right microsaccades as a function of time. Data show group-level mean values. Shadings in panels a and c indicate 95% CI (calculated across 23 participants).

### Aligning EEG data to spontaneous microsaccades reveals transient alpha-power lateralisation

**Figure 1b** shows the time- and frequency-resolved spatial modulation of neural activity (in posterior electrodes PO7/8) aligned to microsaccade onset, according to microsaccade direction. The results reveal a clear transient lateralisation of posterior alpha power (cluster P < 0.001), with relatively lower power contralateral vs. ipsilateral to microsaccade direction. The associated time course of neural lateralisation in the 8-12 Hz alpha band (**Fig. 1c**) confirms how this lateralisation is transient: occurring and disappearing again within 250 ms from microsaccade onset (cluster P < 0.001). Topographical analyses (**Fig. 1d**) revealed the posterior nature of the identified microsaccade-locked alpha lateralisation.

The microsaccade-locked transient lateralisation of alpha activity that we report here is to be distinguished from the previously reported low-frequency increase in power surrounding microsaccades ^15^. As we show in **Supplementary Figure 2**, these two types of microsaccade-locked EEG responses have distinct frequency profiles, distinct topographies, and, most notably, have opposite directions of modulation in the reported lateralisation contrast. In this report, we focus on the alpha-band modulation.

### Similar transient alpha lateralisation for start and return microsaccades

Having uncovered the transient alpha lateralisation surrounding all identified spontaneous microsaccades, we considered whether the transient lateralisation of alpha activity is specific to “start” microsaccades that move away from fixation and therefore are free to go in either direction, or whether it would occur similarly surrounding “return” microsaccades that merely serve to bring gaze back to fixation. To test this, we sorted microsaccades based on their associated displacement of gaze: whether the gaze moved away from (start microsaccades) or back toward (return microsaccades) central fixation.

The gaze-position traces in **Figure 2a** confirm our sorting into start and return microsaccades, separately for left and right microsaccades: start microsaccades resulted in larger gaze deviations from central fixation, while return microsaccades resulted in smaller deviations. Note that microsaccade directions were labelled separately for start and return microsaccades, such that left start microsaccades displace gaze from central fixation to the left, whereas left return microsaccades displace gaze from the right back to the central fixation maker.

**Figure 2.**
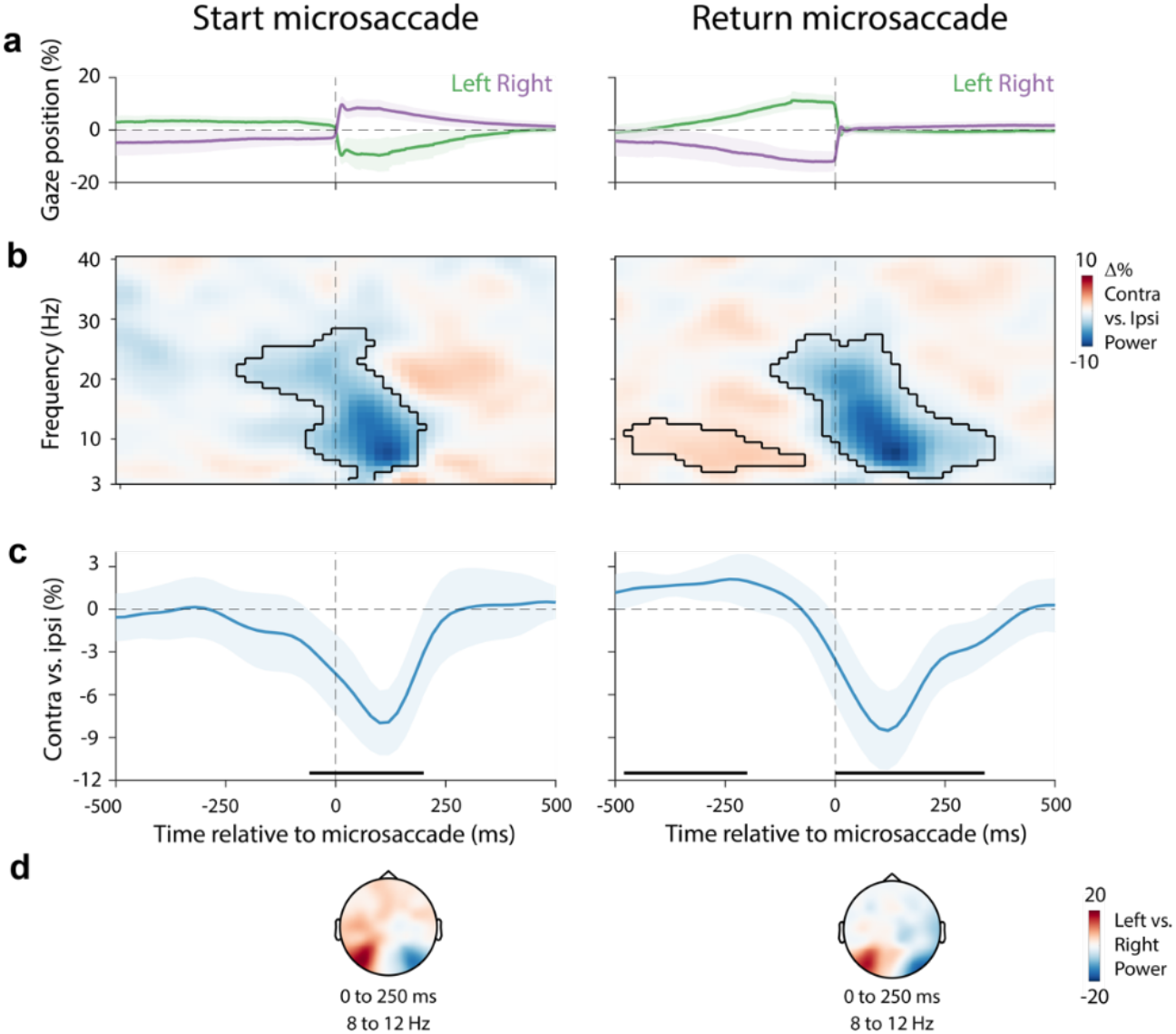
Similar transient alpha lateralisation for start and return microsaccades. **a)** Time course of gaze position for left and right microsaccades for start (left) and return (right) microsaccades. **b)** Spectral lateralisation (contralateral minus ipsilateral) in selected visual electrodes (PO7, PO8) relative to microsaccade direction. Outlines indicate significant time-frequency clusters (two-sided cluster-based permutation test). **c)** Time course of averaged 8-12 Hz lateralisation. Black horizontal lines indicate significant temporal clusters (two-sided cluster-based permutation test). Time courses show mean values, with shading indicating 95% CI (calculated across 23 participants). **d)** Topographical map of the difference between 8-12 Hz power for left vs. right microsaccades.

Our data clearly revealed how microsaccades were associated with transient alpha lateralisation, regardless of whether the microsaccade was away from (**Fig. 2b** left; cluster P < 0.001) or back to (**Fig. 2b** right; cluster P < 0.001) central fixation (**Fig. 2b-d**). In addition, in return-microsaccade trials, we observed the opposite positive cluster (**Fig. 2b** right; cluster P = 0.019) before the onset of microsaccades. This likely reflects the neural modulation associated with the initial start saccade in the opposite direction (increasing confidence in our sorting of these return trials).

### Microsaccade-related alpha lateralisation is driven by an ipsilateral increase in power

We next asked whether the observed alpha lateralisation is driven predominantly by an increase in alpha power ipsilateral to the direction of spontaneous microsaccades or a decrease in alpha power contralateral to microsaccades direction. We focused on start microsaccades because these are less contaminated by preceding microsaccades.

When considering raw power (**Fig. 3a**), we observed a transient increase in the alpha-band signal ipsilateral to microsaccade direction, with no evidence for a contralateral attenuation. To look at this with higher sensitivity, we log-transformed power and subtracted a pre-microsaccade baseline window (−500 to -150 ms relative to microsaccades). This again showed an increase in alpha power ipsilateral to the direction of the microsaccade (**Fig. 3b**), which was particularly evident in the associated time courses of neural lateralisation in the 8-12 Hz alpha band (**Fig. 3c**; cluster P = 0.008). The time course of this ipsilateral alpha-power increase closely matches that of the reported alpha lateralisation in the preceding figures. This suggests that the reported lateralisation is driven by an increase in posterior alpha power ipsilateral to microsaccade direction.

**Figure 3.**
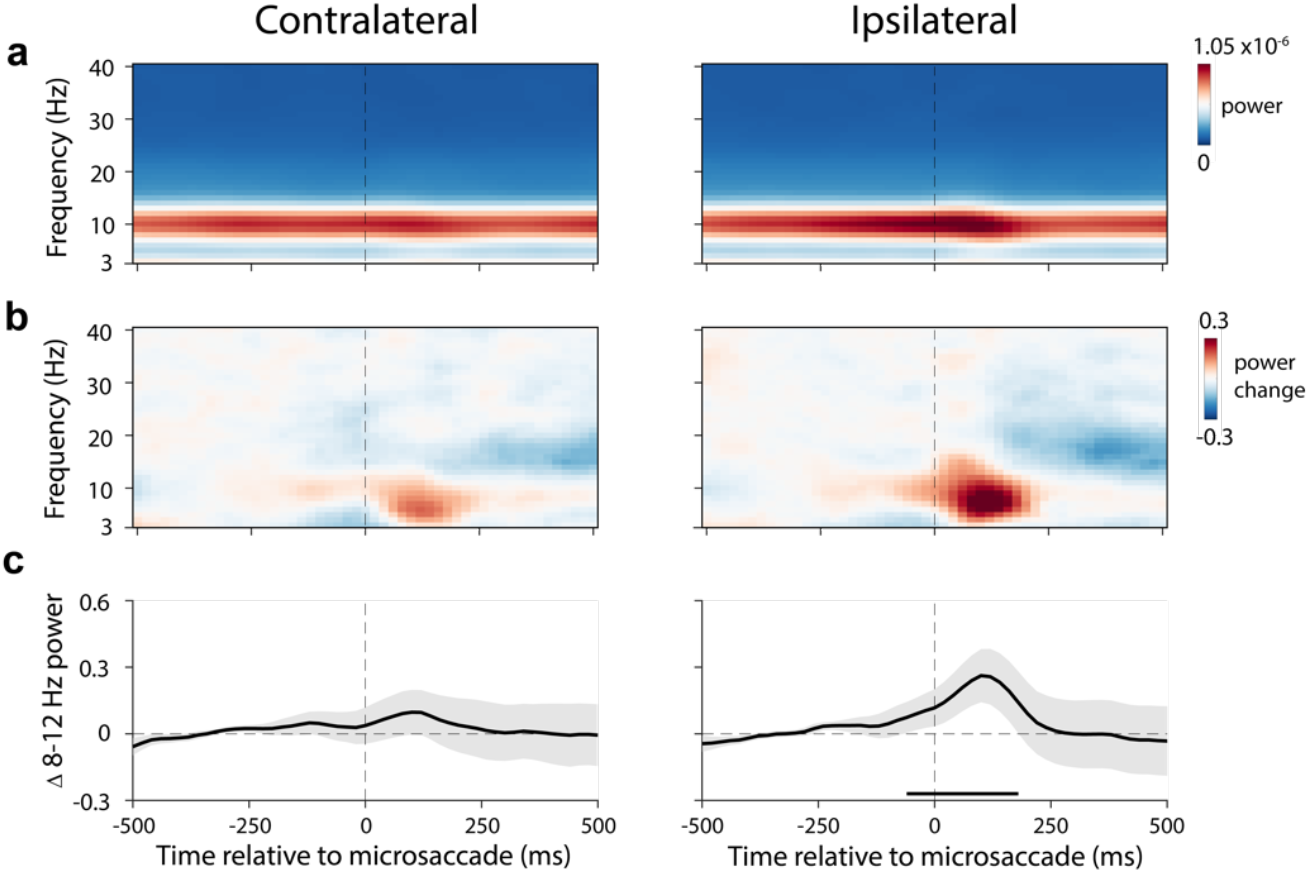
Microsaccades transiently increase alpha power ipsilateral to microsaccade direction. **a)** Time-frequency maps of raw EEG power in selected electrodes (PO7, PO8) contralateral (left) and ipsilateral (right) to microsaccade direction. **b)** Change in log-transformed power, relative to a pre-microsaccade time window (−500 to -150 ms relative to microsaccades). **c)** Time courses of 8-12 Hz power changes corresponding to the data in panel **b**. The black horizontal line indicates the significant temporal cluster (two-sided cluster-based permutation test). Time courses show mean values, with shading indicating 95% CI (calculated across 23 participants).

### The microsaccade-locked increase in ipsilateral alpha power has a phase-locked component

Finally, we asked whether the reported alpha-power modulation has a phase-locked component by considering the inter-trial phase-coherence (ITPC).

As shown in **Figure 4a**, we found a clear increase in low-frequency ITPC in both ipsilateral and contralateral electrodes, consistent with the existence of microsaccade-locked event-related potentials (as documented previously ^15^). Interestingly, when considering microsaccade direction, we found that the ITPC extended to higher frequencies that incorporated the classic 8-12 Hz alpha-band only on the ipsilateral side. A direct comparison between ITPC ipsilateral and contralateral to microsaccade direction converged on a clear difference that was largely confined to the alpha-band (**Fig. 4b**; cluster P < 0.001), occupying a similar time-frequency range as the effect reported in the preceding figures.

**Figure 4.**
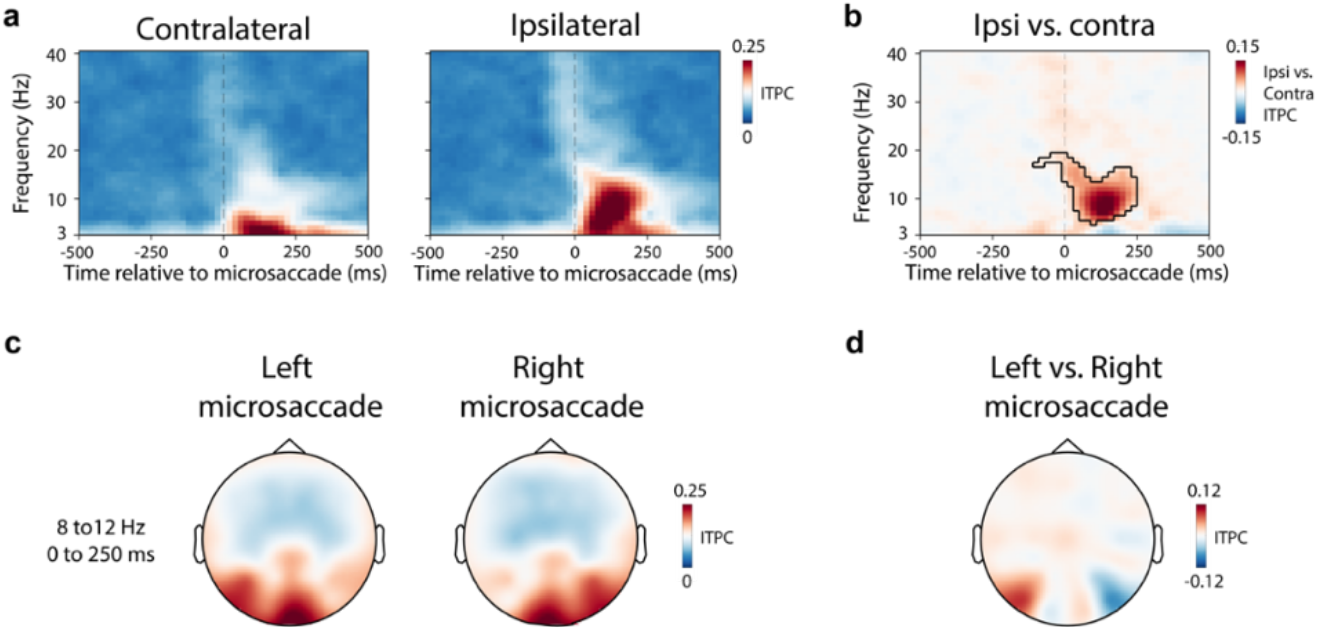
The microsaccade-locked increase in ipsilateral alpha power has a phase-locked component. **a)** Spectral inter-trial-phase-coherence (ITPC) results in selected posterior electrodes (PO7, PO8) contralateral (left) and ipsilateral (right) to microsaccades direction. **b)** The comparison of ITPC between ipsilateral and contralateral electrodes. The outline indicates significant time-frequency clusters (two-sided cluster-based permutation test). **c)** Topographical map of 8-12 Hz ITPC in the 0-250 ms period following left and right microsaccades. **d)** Topography of the ITPC difference following left vs. right microsaccades.

Together, these analyses suggest that the direction of spontaneous microsaccades is associated with a transient lateralisation of posterior EEG alpha power that occurs predominantly ipsilateral to the microsaccade direction, and that has a phase-locked component.

## DISCUSSION

We uncover a transient microsaccade-locked lateralisation of EEG alpha activity according to the direction of microsaccades that were made spontaneously in the absence of any incentive to look to the left or right. This complements work linking lateralised alpha activity to goal-driven (as opposed to spontaneous) saccades ^25–28^ and microsaccades ^10^. It further complements prior work linking microsaccades to non-lateralised M/EEG activity ^14,15,29^ that did not consider microsaccade direction, or that considered lateralised ERPs ^16,17,30^ or behaviour ^31^ rather than spectral alpha modulations in the EEG.

An important open question left unaddressed by the current work is what causes, or mediates, the here-reported link between microsaccade direction and the transient lateralisation of EEG alpha power. We consider at least three possibilities but emphasize that future work is required to disambiguate these scenarios.

First, this link may be mediated by covert spatial attention. Spontaneous microsaccades have been linked to spontaneous shifts of attention ^32–36^, and it is well established that covert attention shifts lateralise EEG-alpha activity ^19–22,37,38^. At the same time, in the delay period that we analysed here, there was no explicit incentive to shift attention: there was nothing to look at to the left or right, and the two visual items in working memory were equally relevant during the delay. Moreover, we found a similar transient alpha lateralisation for start and return microsaccades, showing that the reported effect occurs even for those microsaccades that merely serve to return the gaze to the central fixation marker. Nevertheless, it is conceivable that the observed alpha lateralisation may reflect a consequence of pre-saccadic shifts of attention ^34–36^, which have also been reported surrounding microsaccades ^33,39^.

A second (potentially related) possibility is that spatial alpha modulation by microsaccades serves to inhibit the processing of previously attended (fixated) information, in line with the postulated role of alpha activity in cortical inhibition ^19,20,40^. Consistent with such an interpretation, the observed lateralisation was driven predominantly by posterior sites *ipsilateral* to microsaccade direction that process previously attended parts of space.

Finally, we cannot rule out that mere retinal displacement of the fixation cross may have contributed to (part of) the microsaccade-locked transient lateralisation of alpha activity. It is noteworthy, however, that posterior alpha responses to new visual input (see e.g. ^41–43^) typically peak around 500 ms after stimulation and are sluggish, whereas our microsaccade-locked alpha lateralisation peaked much earlier and was transient. Moreover, even though the retinal displacement by microsaccades is relatively minute in size, the alpha lateralisation reported here was most prominent in relatively lateral electrodes PO7/8 and was distinct from the lower frequency modulations that we observed in more central electrodes O1/2. As such, it is conceivable that the alpha lateralisation reported here is related to prior reports of parafoveal influence of microsaccades at eccentricities far from the retinal displacements associated with the microsaccades themselves ^5,33^.

Our analyses revealed that the reported alpha lateralisation comprised a phase-locked component, consistent with the existence of microsaccade-locked evoked potentials ^15^. Here we not only show how such potentials depend on microsaccade direction but also how they can lead to lateralisation in time-frequency analyses of power where they can appear largely confined to the classical alpha frequency band. We found this to be the case when analysing posterior-lateral EEG electrodes PO7/8 that are commonly used to characterised EEG signals associated with spatial attention and working memory (e.g. ^10,44–48^). Interestingly, the alpha lateralisation complemented low-frequency modulations of power (that are more typically associated with evoked potentials) that we observed in more medial EEG electrodes that had qualitatively distinct profiles (**Supplementary Figures 2-3**).

Whatever the exact physiological origins of the here-reported microsaccade-locked transient alpha lateralisation, our results – together with those before us ^10,14–17,33,49–51^ – carry a relevant practical implication. They highlight the importance of considering microsaccades when reporting neural modulation by specific task variables. This is because conditional differences in microsaccades (such as biases in microsaccade direction ^6–10^) may account for conditional differences in neural activity (such as the spatial modulation of alpha activity ^19–22^) and mislead interpretation. Following this logic, we recently interrogated whether goal-driven biases in microsaccade direction can fully account for the goal-driven lateralisation of EEG-alpha activity by spatial attention ^10^. Like here, we found that alpha lateralisation and microsaccade direction correlated – when microsaccades occurred. However, we also found that microsaccades were not necessary, since task-driven alpha lateralisation was preserved in trials without microsaccades ^10^ (see also ^52^). The present report suggests that explorations of this type should be adopted more widely. Simply assuming that microsaccades – because of their small size – are unlikely to contribute to specific task-related neural modulations may no longer be an acceptable premise.

## METHODS

The current results present the re-analysis of a dataset from an experiment in which human participants performed a working-memory task while electroencephalography (EEG) and eye-tracking data were recorded concurrently. Complementary results from this dataset have been reported in prior studies, addressing distinct questions ^9,10,53^. Unlike all prior reported analyses from this dataset, we here focused on microsaccades during the delay period of a working-memory task when only a fixation marker remained on the screen that participants were instructed to fixate. By aligning EEG data to these microsaccades in the delay period, we aimed to investigate the association between spontaneous microsaccades (made in the absence of any incentive to look elsewhere but at the central fixation marker) and the lateralisation of EEG-alpha activity according to microsaccade direction.

### Participants

Twenty-five healthy human participants voluntarily participated in the study (age range 19-36; 11 male, 2 left-handed). Participants were compensated 15 GBP per hour for their time. The sample size was based on previous publications from the lab that had similar designs and focused on similar neural signatures (for example see ^54^). As in our two prior articles that also focused on the eye-tracking data from this dataset (Experiment 1 in ^9^ and ^10^), two participants were excluded due to their poor quality of eye-tracking data. The experimental procedures were reviewed and approved by the Central University Research Ethics Committee of the University of Oxford.

### Task and procedure

Participants performed a working-memory task in which they were required to encode and maintain two visual items (tilted coloured bars) in working memory and later report one of them. Each trial began with a brief (250 ms) encoding display that contained two to-be-memorised bars with different colours and orientations. The encoding display was followed by a retention delay in which only a central fixation cross remained on the screen for a variable delay time that was randomly drawn between 2,000 and 2,500 ms. Participants were instructed to keep fixation during this retention delay and had no incentive to look elsewhere. The results reported here exclusively concern the data from this delay period, during which participants attentively fixated while awaiting a colour change of the central fixation cross that instructed them about which memory item to report. For this current report, the most important aspect of our task was that participants engaged in a period of attentive fixation, with no incentive to look elsewhere than at the fixation cross. We report other aspects of our task for completeness, even if they may not be critical for the findings we present in the current report.

After the delay, the central fixation cross changed colour to match one of the two memory items, designating it as the target for the report. At this stage, participants were required to report the orientation of the colour-matching memory item by using a keyboard to adjust a central reporting dial. The reporting dial was always presented centrally (surrounding the fixation cross), regardless of whether the left or right memory item was probed for the report.

In the encoding display, the two bars (size: 5.7° ×0.8°) were centred at approximately 5.7° to the left and right of fixation. In each trial, the bars were randomly allocated two distinct colours out of four possible colours: blue (RGB: 21, 165, 234), orange (RGB: 234, 74, 21), green (RGB: 133, 194, 18) and purple (RGB: 197, 21, 234), and two distinct orientations (one left tilted and one right tilted) between ±20° to ±70°. Colours were drawn independently of bar location and tilt across trials. At the end of the trial, the report of the target orientation was achieved by pressing and holding down one of two keys on the keyboard (the ‘\’ key to rotate the dial leftward and the ‘/’ key to rotate the dial rightward) and the response was terminated at key release. The reporting dial always started in the vertical position at response initiation. After response termination, feedback was immediately given by changing the colour of the fixation cross to green for 200 ms for reproduction errors less than 20°, and to red otherwise. After the feedback, an inter-trial interval (randomly drawn between 500 and 800 ms) was introduced before the next encoding display.

After practising the task for 5-10 minutes, participants completed two consecutive sessions of one hour each, with a 15-minute break in between. Each session contained 10 blocks of 60 trials, yielding 1,200 trials per participant.

The experiment was controlled by Presentation (version 18.3, Neurobehavioral Systems Inc., Berkeley, CA). During the experiment, participants sat in front of a monitor (100-Hz refresh rate) at a viewing distance of approximately 95 cm with their heads resting on a chin rest.

### Eye-tracking acquisition and pre-processing

The eye tracker (EyeLink 1000, SR Research, sampling rate = 1000) was positioned on the table ∼15 cm in front of the monitor and ∼80 cm away from the eyes. The gaze positions along the horizontal and vertical axes were continuously recorded for both eyes. Before recording, the built-in calibration and validation protocols from the EyeLink software were used to calibrate the eye tracker.

After recording, the eye-tracking data were firstly converted from the original .edf to the .asc format. They were then read into MATLAB using the Fieldtrip toolbox ^55^, which we also used for EEG analyses. By merging the data from the left and right eyes, we kept a single horizontal and a single vertical gaze-position channel for further analysis. The blinks were marked by detecting NaN clusters in the eye-tracking data. Then, all data from 100 ms before to 100 ms after the detected NaN clusters were also set to NaN to eliminate residual blink artefacts.

### Microsaccade detection

Our focus in the current study was on EEG-alpha lateralisation that is most well characterized along the horizontal (left/right) axis. Accordingly, our eye-tracking analysis focused on the horizontal channel of the merged eye data to be optimally sensitive to detecting left vs. right shifts of gaze. To identify microsaccades, we employed a velocity-based method that we developed and validated in our prior study that focused on the relation between attention-driven microsaccades and alpha activity after the attentional cue ^10^. Here, we instead focused on ‘spontaneous microsaccades’ that occurred during the delay period.

In short, gaze velocity was calculated using the absolute value of the temporal derivative of gaze position. Then velocity was smoothed with a Gaussian-weighted moving average filter with a 7-ms sliding window (using the built-in function “smoothdata” in MATLAB). The samples in which the velocity exceeded a trial-based threshold of 5 times the median velocity were marked as the onset of a saccade. Furthermore, to avoid counting the same eye movement multiple times, a minimum delay of 100 ms between successive gaze shifts was imposed. Saccade magnitude and direction were calculated by estimating the difference between pre-saccade gaze position (−50 to 0 ms before saccade onset) vs. the post-saccade gaze position (50 to 100 ms after saccade onset). To zoom in on microsaccades, we only used the detected saccades whose magnitude was < 1 visual degree (which was the case for the majority of our detected gaze shifts during the delay; **Supplementary Figure 1**). We also excluded microsaccades with horizontal displacement smaller than 1% toward the stimuli, corresponding to 3.42 min of arc, which is comparable to exclusion criteria in other microsaccade studies (e.g. ^56–58^).

In addition to detecting microsaccades and defining their direction as leftward or rightward, we aimed to separate microsaccades by whether they moved away from (“start microsaccade”) or moved back toward (“return microsaccade”) fixation. To this end, we additionally estimated the pre- and post-saccade distance from the fixation position. If the post-saccade distance was larger than the pre-saccade distance, we defined the saccade as a start microsaccade. If the post-saccade distance was smaller, we defined the saccade as return microsaccade. To minimise the contribution of gaze drift, for this analysis we estimated gaze positions associated with central fixation as the median gaze position in the larger fixation time window of interest (from -1.5 to -0.5 ms relative to cue onset).

For the current analyses, we focused exclusively on microsaccades that occurred during the delay period, while ensuring at least 500 ms after the offset of the visual encoding display, and at least 500 ms prior to cue onset at the end of the delay. All microsaccades detected in this period were defined as ‘spontaneous’, provided there was no incentive to look anywhere else but at the fixation cross.

### EEG acquisition and basic processing

EEG data were collected using Synamps amplifiers and Neuroscan acquisition software (Compumedics) at a sampling rate of 1000 Hz. 61 electrodes were distributed across the scalp according to the international 10–10 system. Vertical and horizontal EOG were measured to monitor ocular artefacts (to guide their later removal using ICA). Data were referenced to the left mastoid and filtered between 0.1 and 200 Hz during acquisition. After the acquisition, data were re-referenced to the mean of the right and left mastoids.

EEG data were analysed in Matlab, using a combination of Fieldtrip ^55^ and custom code. Independent Component Analysis (ICA), as implemented in Fieldtrip, was performed to identify and remove the components associated with eye blinks and eye movements (with bad components identified based on their correlation with the measured EOG signals). After ICA correction, trials with exceptionally high EEG variance were identified and excluded using the “ft_rejectvisual.m” function in Fieldtrip with the ‘summary’ method. Finally, to increase the spatial resolution, a surface Laplacian transformation was performed.

### Alignment of EEG and microsaccades

To characterize the EEG signal by microsaccades, we aligned all EEG data to the onset of the detected spontaneous microsaccades within the pre-defined delay period of interest (including only microsaccades that occurred at least 500 ms after visual encoding offset and the latest 500 ms prior to cue onset at the end of the working memory delay period). The EEG data were epoched from 1200 ms before to 1200 ms after the onset of microsaccades. This resulted in a total of 27167 epochs (1181 M ± 106 SEM epochs per person) for our later analysis.

### Electrode selection

In the current experiment, we focused on the left/right posterior PO7/8 electrodes. These are commonly used to characterise spatial modulations of EEG-alpha activity (for example during visual-spatial attention and visual working memory; e.g. ^10,44,45,47,48,59^) and were also used in our prior reports ^10,53^. In addition, we considered EEG activity in more medial left/right posterior electrodes O1/2, which have previously been used to study microsaccade-locked event-related potentials ^15^.

### Time-frequency analysis

The microsaccade-aligned EEG time series were decomposed into a time-frequency representation using a short-time Fourier transform of Hanning-tapered data, as implemented in Fieldtrip. The spectral power and phase between 1 and 50 Hz in 1-Hz steps were estimated using a 300-ms sliding time window that was advanced over the data in steps of 20 ms.

To estimate the lateralised modulation of spectral power relative to microsaccade direction (left/right), we expressed this lateralisation as normalised contrasts (i.e., ((contra-ipsi)/(contra+ipsi)) × 100) and then averaged these contrasts across the predefined left and right electrodes. In addition, we estimated the time courses of lateralisation by averaging over all frequencies within the pre-defined and well-established 8-12 Hz alpha band of interest. Finally, for completeness, we visualised topographical maps by contrasting left and right microsaccade trials (i.e., ((left-right)/(left+right)) × 100) for all electrodes.

In addition to the direct quantification of spectral lateralisation, we aimed to separately estimate the power change in electrodes ipsilateral and contralateral to the direction of microsaccades. To this end, we normalized power using the log-transform (using the built-in function “log” in MATLAB) and baseline-corrected the contralateral and ipsilateral data by subtracting the pre-microsaccade log-normalised power in the baseline window from -500 to -150 ms relative to microsaccade onset.

Complementary to the main analyses of spectral power, we also considered inter-trial phase coherence (ITPC ^60,61^). First, Fourier coefficients were normalized to vector-length one by dividing the complex coefficients by their absolute vector length. We then averaged the length-normalized complex coefficients across all trials (separately for each time- and frequency point) and calculated ITPC as the vector-length of the resulting trial-averaged two-dimensional coefficient. The ITPC value represents a measure of phase consistency across trials ranging from 0 to 1, with 1 reflecting perfect phase consistency across trials.

### Statistical analysis

We used a cluster-based permutation approach ^62^ to statistically evaluate our time-frequency data of interest. This well-established cluster-based permutation approach enabled us to evaluate the reliability of neural patterns at multiple neighbouring data points (i.e., across time and frequency) while circumventing the multiple-comparison problem. Clusters were defined following default settings in Fieldtrip (identifying time-frequency points in which univariate two-sided paired samples t-test was significant and clustering neighbouring samples that met this criterion, then defining the size of a cluster as the sum of all t values in the cluster). After identifying clusters in the original data, we used 10,000 random permutations of the trial-averaged data to create a permutation distribution of the largest clusters that were found after each permutation. The P value for the clusters observed in the original data was defined as the proportion of random permutations that yielded a larger cluster.

## Data availability

All data used for this report have already been made publicly available through the Dryad Digital Repository. The eye-tracking data can be found at: https://doi.org/10.5061/dryad.m99r286 ^63^ (Experiment 1). The corresponding EEG data can be found at: https://doi.org/10.5061/dryad.sk8rb66 ^64^.

## Code availability

Relevant code associated with the here-presented analyses will be made available through GitHub before publication.

## Supplementary Information – Figures S1-S3

**Supplementary Figure 1.**
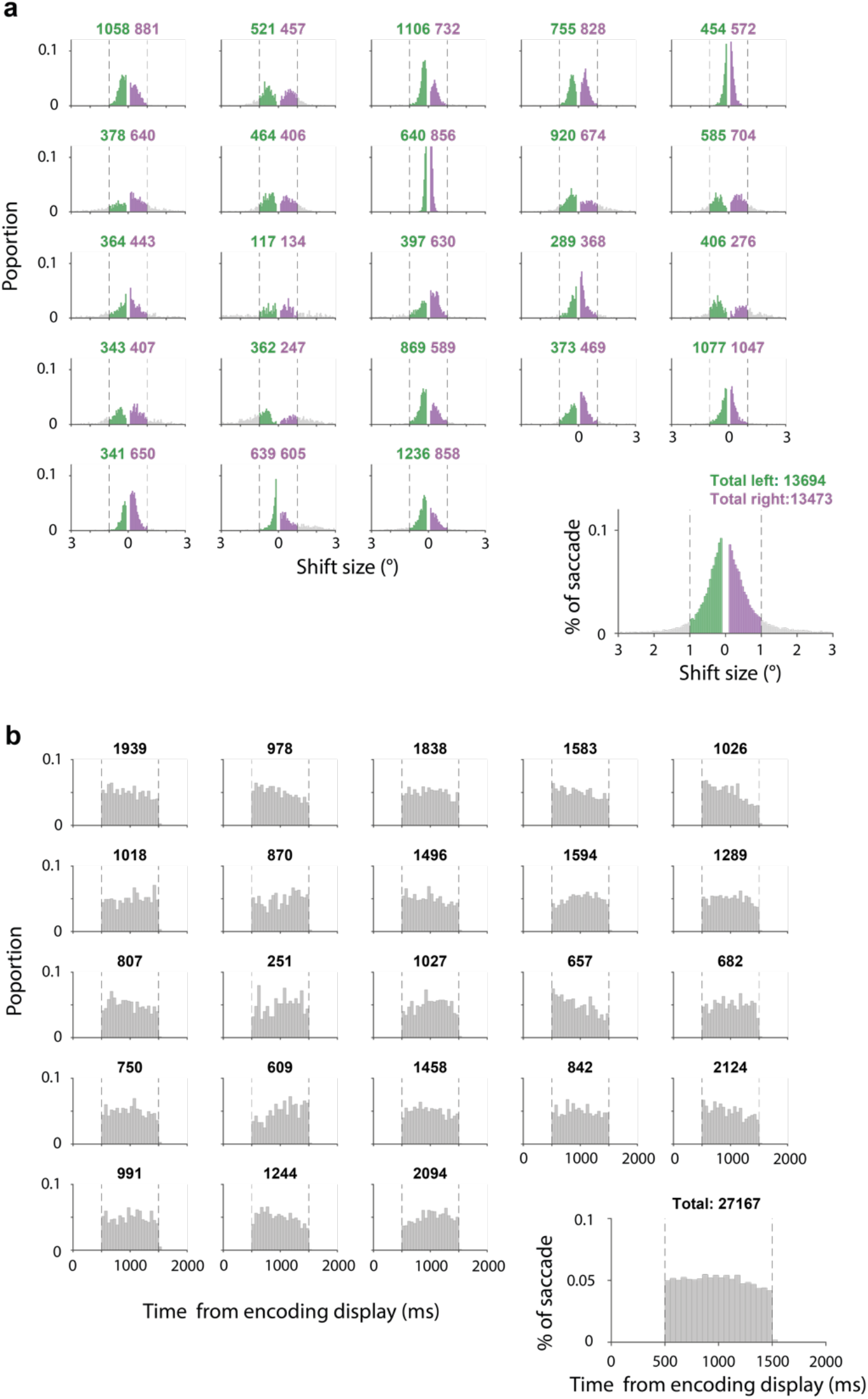
Spatial and temporal distributions of identified microsaccades across participants. **a)** Direction and magnitude distributions of detected horizontal saccades during the retention delay. The dashed line represents the threshold (1-degree visual angle) under which the detected saccades were treated as ‘microsaccades’ and used in the reported analysis. **b)** The temporal distribution of the identified usable microsaccades across the delay period. In both panels, each plot represents an individual participant, while the right bottom plot represents the distribution of detected microsaccades aggregated across all participants.

**Supplementary Figure 2.**
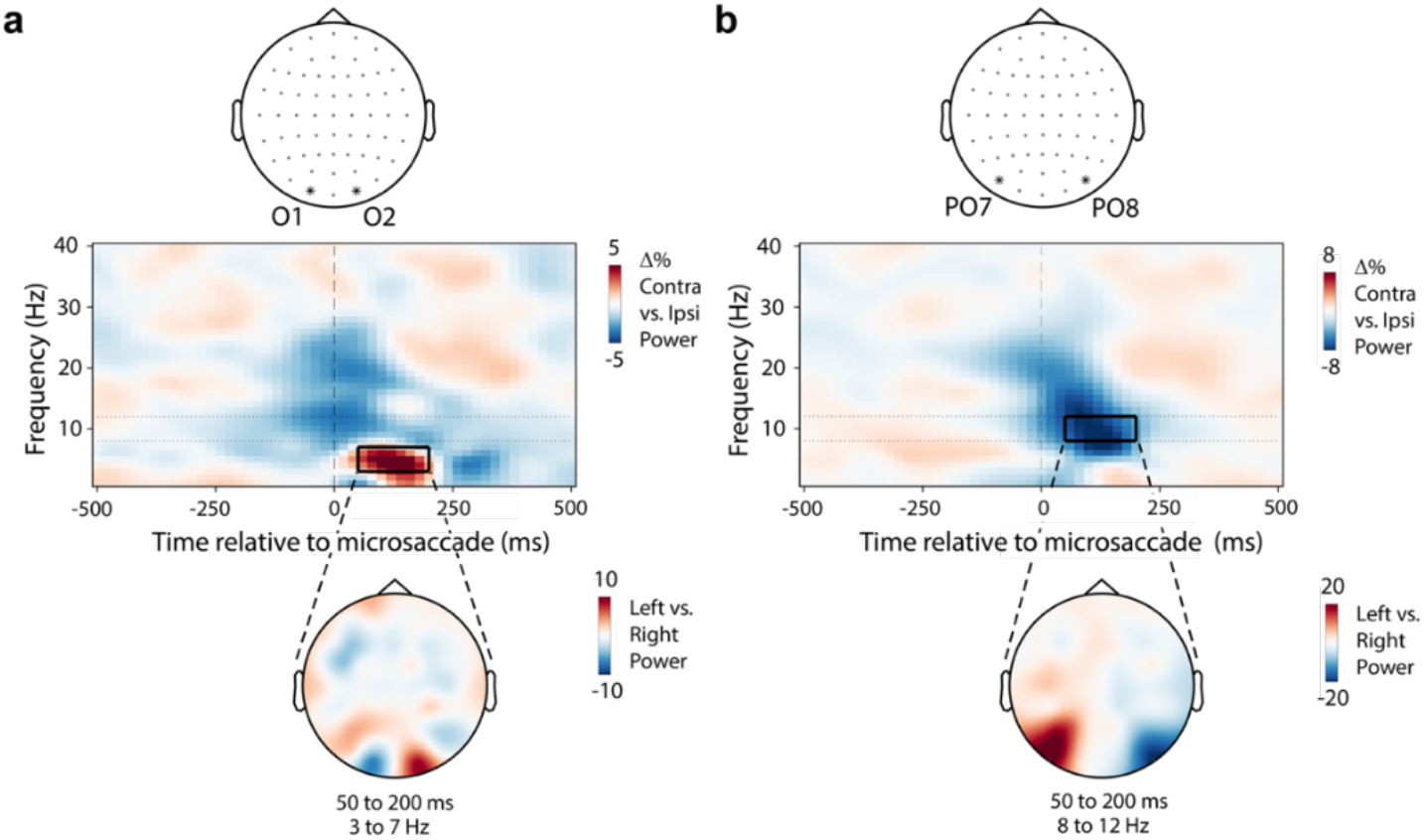
Microsaccade-locked EEG responses in 3-7 Hz and 8-12 Hz have distinct topographies and opposite directions of modulation. **a)** Analyses like in **Main Figure 1**, but instead of using electrodes PO7/8, we used electrodes O1/O2. **b**) equivalent results using electrodes PO7/8 instead for reference (same data as in **Main Figure 1**).

**Supplementary Figure 3.**
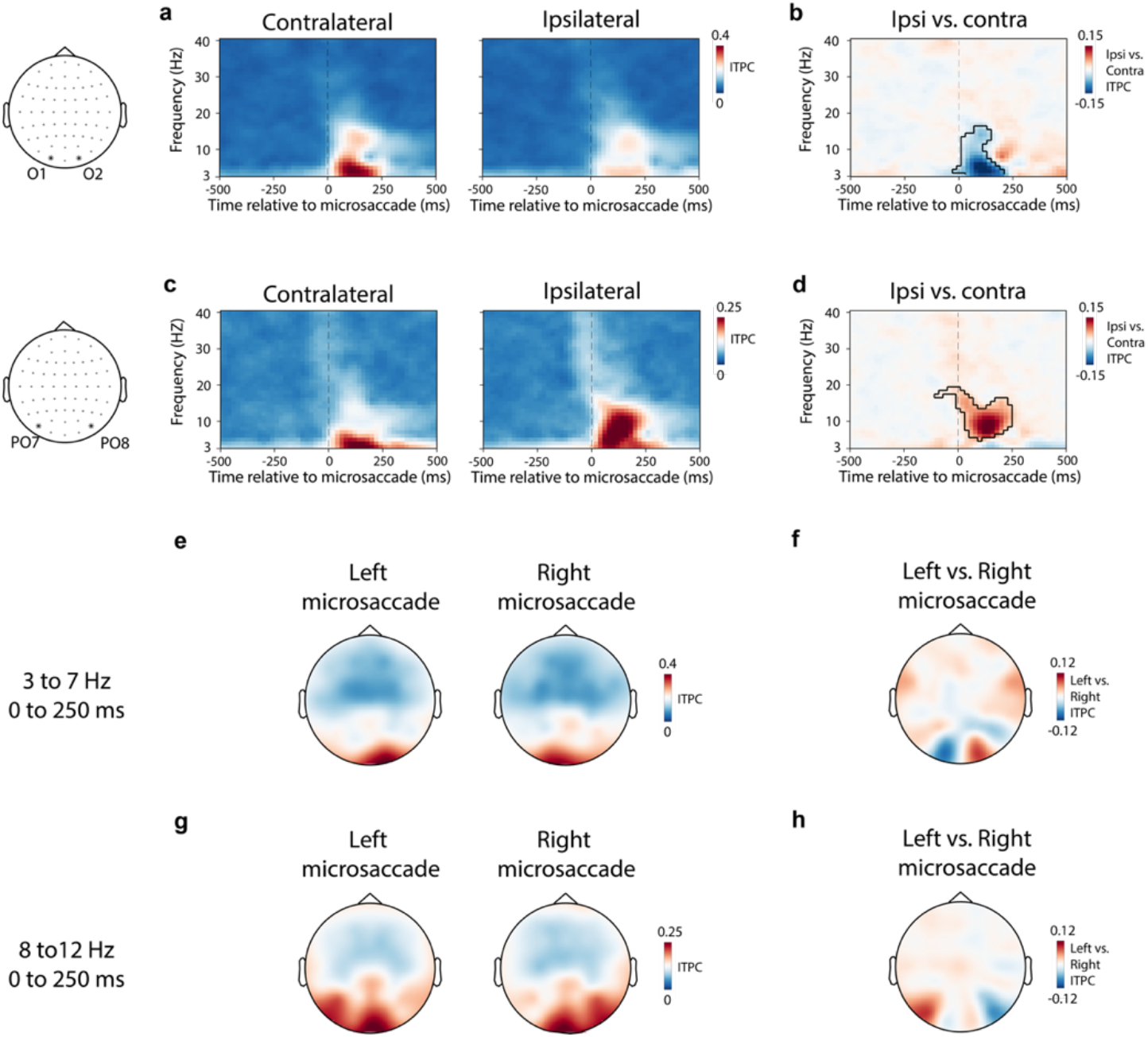
Inter-trial phase-coherence as a function of electrodes (O1/2 and PO7/8). Conventions as in **Main Figure 4**, separately for electrodes O1/2 (a-b) or PO7/8 (c-d) and for 3-7 Hz (e-f) and 8-12 Hz (g-h).

